# Cryo-EM structure of DNA polymerase θ helicase domain in complex with inhibitor novobiocin

**DOI:** 10.1101/2023.01.20.524915

**Authors:** Hanbo Guo, YanXia Wang, Jun Mao, Huimin Zhao, Yuntong He, Yuandong Hu, Jing Li, Yujie Liu, Zheng Guan, Allen Guo, Xiaodan Ni, Fengying Zhang, Jie Heng

## Abstract

DNA double-strand breaks (DSBs) are highly toxic lesions that occur during the cellular metabolic process. DNA Polymerase theta (Polθ) is an error-prone polymerase that has been implicated in the repair of chromosome breaks, recovery of broken replication forks, and translesion synthesis. The inhibition of Polθ activity has been implicated in killing HR-deficient tumor cells *in vitro* and *in vivo*. We present the first biochemical evidence that the antibiotics novobiocin (NVB) noncompetitively inhibit ATP hydrolysis by the ATPase domain of the Polθ helicase domain (Polθ-HLD). We report the Cryo-EM structure of apo dimeric Polθ helicase domain (Polθ-HLD), and the first inhibitor occupied Polθ-HLD structure. Our structure identifies a non-canonical novobiocin binding pocket, distinct from the canonical site that partially overlaps with the ATP in the ATPase domain. Comparison with the homolog helicase Hel308-DNA duplex complex suggests that the novobiocin competitively binds to a triangle hub on the DNA translocation pathway and blocks the ssDNA binding and translocation. Furthermore, the first dimeric structure of Polθ-HLD also provides a structural framework for revealing the microhomology-mediated end-joining mechanism. Our results demonstrate that the inhibitor-occupied structure combined with rational, structure-based drug design will undoubtedly accelerate the discovery of potent inhibitors with better efficacy and target selectivity to human Polθ.

## Introduction

Stochastic DNA double-strand breaks (DSBs) are one of the most deleterious types of DNA lesions in eukaryotic cells^1^. The inability to repair properly to DNA damage may lead to genetic instability and cell death, which in turn may enhance the rate of cancer development^2,3^. There are two distinct and complementary mechanisms for DNA DSB repair: homologous recombination (HR)^4^ and canonical non-homologous end-joining (NHEJ)^5^. Nevertheless, many cancer types that are deficient in HR- and NHEJ-dependent proteins can rely on a third DSB repair pathway, DNA polymerase theta (Polθ)-mediated end joining (TMEJ), which is an alternative error-prone DSB repair pathway that uses sequence microhomology to recombine broken DNA ends^6^. Accordingly, Polθ is synthetic lethal with a number of genes frequently mutated in cancer, including HR factors^7^ and DNA damage response genes^8^. Developing first-in-class Polθ-targeting inhibitors in combination with other synthetic lethal inhibitors represents a novel therapeutic strategy for cancer treatment.

Polθ is a multifunctional enzyme that contains an N-terminal conserved superfamily 2 helicase domain (Polθ-HLD), an unstructured central region, and a C-terminal A-family DNA polymerase domain (Polθ-POL)^9^. Polθ can bind to long single-stranded DNA (ssDNA) overhangs generated by 5′–3′ resection of DSBs and anneals sequences with 2–6 base pairs of microhomology to use them as primers for DNA synthesis^10^. The structure-function analyses reveal that the Polθ-HLD not only has classical ATPase and helicase activity but also competes with HR factors, like RPA^11^ and Rad51^7^, for resected single-stranded DNA-ends, and the Polθ-POL is responsible for DNA synthesis either using its terminal transferase or templated extension activity. The available crystal structure of both helicase domain^12^ and polymerase domain^13^ in combined with *in vitro*^10,14^ and *in vivo*^15^ biochemical results provide a comprehensive landscape for analyzing the mechanistic multifunction of the Polθ. Recently, selective inhibitors, ART558^16^ and RP-6685^17^, targeting the Polθ-POL, is reported and suggested as promising candidates in synthetic lethality-based anticancer therapy. Although the helicase domain is also proved indispensable for the translesion synthesis and microhomology-mediated end-joining (MMEJ) of long ssDNA overhangs, inhibitors targeting this domain are less reported, except an antibiotic novobiocin (NVB)^18^, which was suggested to inhibit the ATPase activity and phenocopy Polθ depletion specifically. Nevertheless, the structural basis of the NVB binding pocket and the precise molecular mechanism of how NVB modulates Polθ-HLD activity remain compelling questions.

Here, we report the Cryo-EM structure of dimeric Polθ-HLD in the apo and in complex with inhibitor NVB. The NVB, a well-known aminocoumarin antibiotic that inhibits the ATPase activity of Polθ-HLD with a half-maximum inhibitory concentration of 24 μM, was predicted to bind to a tunnel within the Polθ ATPase domain through molecular docking. Inconsistent with the docking model, our structure reveals that the NVB binds to a non-canonical binding pocket formed by domains 1, 2, and 4, which is distinct from the canonical binding site within the ATPase subunit. The coumarin core moiety of NVB plays a hub role in stabilizing the interface formed by the ratchet domain and the ATPase domain, which suggests an allosteric modulation mechanism to the ATPase activity. Since the Polθ exhibits ssDNA-dependent ATPase activity, it was suggested that ssDNA regulates the ATPase activity of Polθ by positively allosteric modulation^9,19^. Comparison with the homolog helicase Hel308-DNA complex^20^ indicates that the NVB competitively binds to a triangle hub on the DNA translocation pathway and blocks the ssDNA binding and translocation. Moreover, our Cryo-EM structure validates that Polθ-HLD exhibits dimer in the solution other than the tetramer observed in the crystal structures. The dimerization of Polθ-HLD also provides a structural framework for understanding the microhomology-mediated end-joining mechanism. Overall, our results provide structural insight into the inhibitory mechanisms of Polθ activity, which will accelerate the discovery of analog inhibitors with better affinity, efficacy, and selectivity for clinical cancer treatment.

## Result

### The validation of the inhibitory role of different compounds to Polθ-HLD

In line with the available evidence that Polθ can act on 3’ ssDNA overhangs to promote MMEJ of DSBs repair^21^, the ssDNA overhang was suggested to require the Polθ-HLD attachment in a recombinant system with purified full-length Polθ^22^. Meanwhile, the purified human Polθ-HLD was reported to bind to ssDNA relatively tightly compared with different types of DNA structures. Therefore, the ATPase activity of Polθ-HLD can be allosterically modulated by ssDNA^19^. Hence, we first set up the ATPase activity assay to evaluate the functionality of purified Polθ-HLD by following a well-established ADP-Glo luminescent method. The result shows that ssDNA stimulates the ATPase activity in a concentration-dependent manner with an EC50 of about 18.3 nM (Figure S1.a). Then, we test the inhibitor effect of NVB on ATPase activity, which displays an IC50 of about 13uM (Figure S1.b), similar to the reported value of 24uM^18^. Interestingly, the enzyme inhibition analysis with different NVB concentrations indicates that the NVB is a noncompetitive inhibitor to the ATPase activity (Figure S1.c), which contradicts the hypothesis that the NVB binds near the canonical ATP binding site in the ATPase subunit, like DNA gyrase^23^. Furthermore, additional results suggest that the NVB competitively inhibit the ssDNA binding to the Polθ-HLD (Figure S1.d).

### The overall structure of the Polθ-HLD in apo and in complex with NVB

We determine the Cryo-EM structure of the human Polθ-HLD in the apo state at 3.27 Å and in complex with inhibitor NVB at 3.14 Å (Figure 1.a-c). The overall structures of the apo state and the inhibitor binding state are very similar, with an average root mean square deviation (r.m.s.d) value of 0.94 Å. Notably, in the Cryo-EM structures, the Polθ-HLD forms a homodimer other than a tetramer observed in the crystal structures^12^. It was consistent with the hypothesis that dimer may be the minimum unit of this enzyme, and two Polθ protomers can work together to join either side of a DNA break to facilitate the microhomology annealing step^24^. A detailed summary of the Cryo-EM data collection, refinement, and validation statistics is given in Table 1.

**Table 1.** Cryo-EM data collection, refinement, and validation statistics.

|  | Apo Polθ-HLD<br>Consensus map | Polθ-HLD<br>Consensus map |
| --- | --- | --- |
| **Data collection and processing** |  |  |
| Microscope | KriosG4 | Titan KriosG3i |
| Camera | Falcon 4 | K3 |
| Imaging mode | Counted resolution | Counted super resolution |
| Magnification | 96k | 105k |
| Voltage (kV) | 300 | 300 |
| Electron exposure (e<sup>-</sup>/Å<sup>2</sup>) | 51.07 | 50 |
| Defocus range (μm) | -1.0~-2.0 | -1.0~-2.0 |
| Pixel size (Å) | 0.86 | 0.669 |
| Symmetry imposed | C1 | C1 |
| Initial particle images (no.) | 451,012 | 655,791 |
| Final particle images (no.) | 265,025 | 311,924 |
| Map resolution (Å) | 3.27 | 3.14 |
| FSC threshold | 0.143 | 0.143 |
| Map resolution range (Å) | 2.0-5.0 | 2.5-4.5 |
| **Refinement** |  |  |
| Initial model used (PDB code) | AF2 predicted | AF2 predicted |
| Model resolution (Å) | 3.27 | 3.14 |
| FSC threshold | 0.143 | 0.143 |
| Model resolution range (Å) | 2.0-5.0 | 2.5-4.5 |
| Map sharpening *B* factor (Å<sup>2</sup>) | -137.2 | -139.2 |
| Model composition |  |  |
| Protein residues | 1510 | 1395 |
| Ligands | 0 | 2 |
| R.m.s. deviations |  |  |
| Bond lengths (Å) | 0.003 | 0.003 |
| Bond angles (°) | 0.562 | 0.570 |
| Validation |  |  |
| Clashscore | 7.09 | 8.97 |
| Rotamers outliers (%) | 0.00 | 0.00 |
| Ramachandran plot |  |  |
| Favored (%) | 97.61 | 97.06 |
| Allowed (%) | 2.39 | 2.87 |
| Disallowed (%) | 0.00 | 0.07 |

**Fig. 1.**
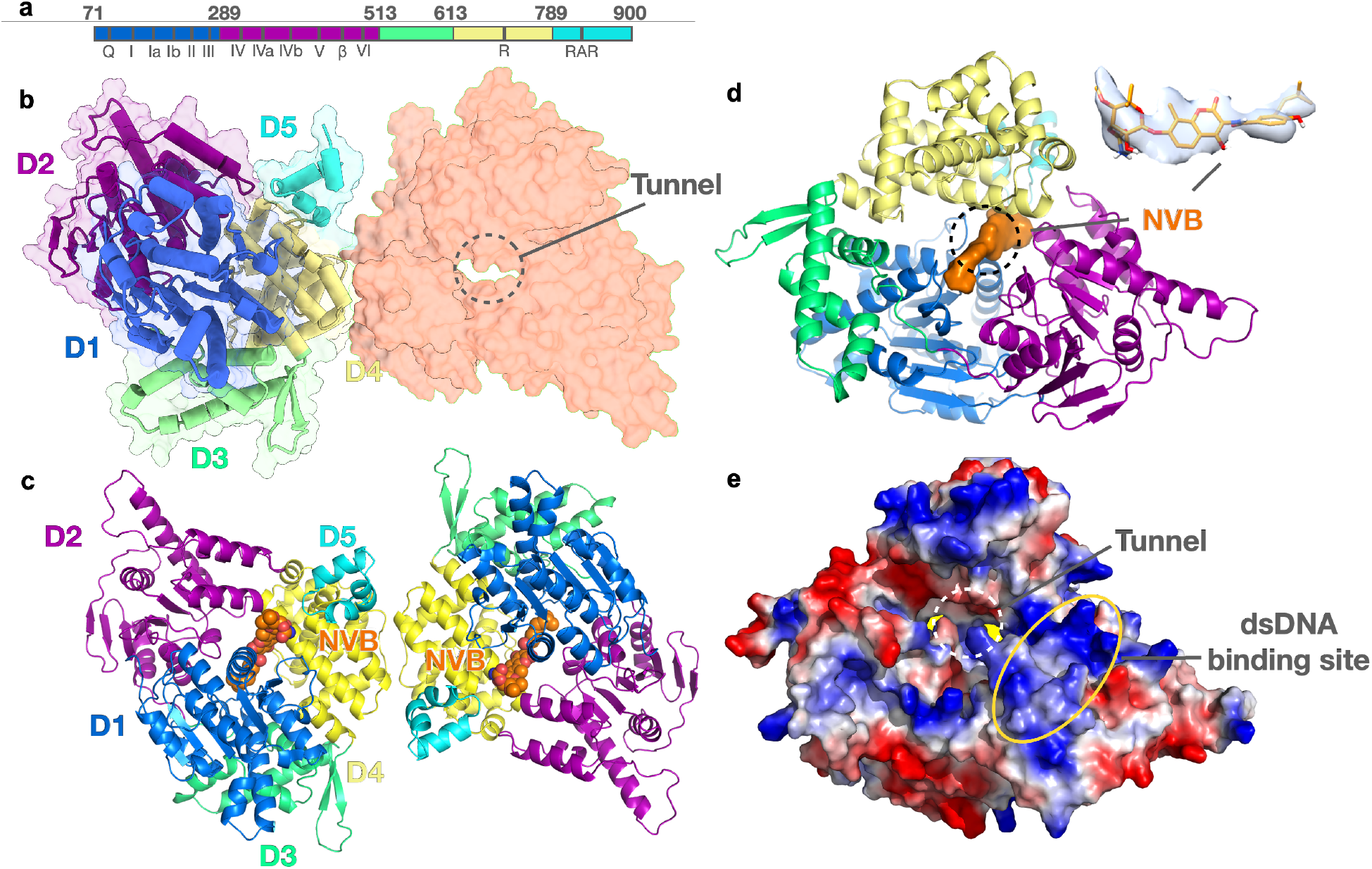
The overall structure of dimeric Polθ-HLD in complex with novobiocin. (a). Schematic domains and conserved motifs of Polθ-HLD. Domain boundaries are indicated on top, and sequence motifs beneath with abbreviations and roman numerals. Q: motif Q; β: β-hairpin; R: ratchet helix; RAR: RAR motif. Domains 1 to 5 are colored blue, purple, green, yellow, and cyan respectively. (b). The apo dimeric Polθ-HLD, NVB is shown as brown spheres. All five domains pack together to form a ring-shaped structure with a central tunnel for ssDNA binding. (c). The dimeric Polθ-HLD-NVB complex, NVB is shown as brown spheres. (d). The ligand density of NVB locates in the central tunnel. (e). The electrostatic surface of Polθ-HLD, the putative dsDNA binding site is highlighted with a yellow ellipse.

As previously defined^12^, the structure of Polθ-HLD consists of five structural domains (domains 1-5, D1-D5). The N-terminal region of Polθ-HLD is composed of two tandem RecA-like domains (D1 and D2), which consist of several characteristic sequence motifs that are important for ATP and DNA binding, named motif Q, I, Ia, Ib, and II to VI (Figure 1.a, and Supplementary Figure S2). The middle region is a winged helix domain (D3), which plays a role as a hinge to permit tightly binding around the DNA substrate. The fourth and fifth domains are the ratchet domain (D4), which coordinates the DNA translocation and ATPase function, and the helix-hairpin-helix domain (D5), which interact with the 3’ ssDNA tail. All five domains pack together to assemble a ring-shaped structure (Figure 1.b) as other structural homologs, including archaeal helicases Hel308^20,25^ and Hjm^26^. The central polar tunnel of Polθ-HLD was wrapped by domains 1, 2, and 4 (Figure 1.b).

We incubate a 50-fold excess molar ratio of NVB with apo Polθ-HLD at a protein concentration of 1 mg/mL, and subsequently vitrified for Cryo-EM study. By comparing with the apo structure, we identify a strip density of NVB in the central tunnel in the NVB-Polθ-HLD complex structure (Figure 1.c,d). The binding site is in a cleft formed by D1, D2, and D4, which supports the observation that NVB increases the Polθ stability in a dose-dependent manner^18^. Furthermore, the extensive positively charged surface inside and outside the central tunnels, especially the surface from D2 (Figure 1.e), implies a potential nucleic acid binding interface supported by the reported crystal structure of DNA duplex-Hel308 complex^27^.

### The inhibitor NVB binding sites

Novobiocin is an aminocoumarin antibiotic initially approved in 1964 for the treatment of severe infections due to susceptible strains of *Staphylococcus aureus*. Since the NVB functioned as a competitive inhibitor of the ATPase reaction catalyzed by the GyrB subunit of the bacterial DNA gyrase enzyme, it was suggested that the NVB molecule partially overlapped with the ATP binding site^23,28^. The canonical NVB binding site is next to the ATP binding site, with the novobiose sugar overlap with the adenine ring of ATP. Besides, NVB also binds to other types of target proteins, including topoisomerase IV (ParE), which is essential for chromosome segregation^30,32^, heat shock protein 90 (HSP90)^33^, autophagy-related protein LC3A^36^, and lipopolysaccharide (LPS)-transport proteins^34,35^. In the case of the LPS transporter, the novobiocin binds to a non-canonical allosteric modulation site away from the ATP binding site.

Although the preliminary docking result indicates that NVB bind near the ATP binding site, the non-competitive evidence from biochemical assay support that the NVB may bind to a distinct non-canonical site away from the canonical site in the ATPase domain. Residues form the NVB pocket from the D1, D2, and D4, including conserved Motif Ia, Motif Ib (part 1), Motif Ib (part 2), Motif II, Motif IVa, and the Ratchet helix (Figure 2.a,b). Notably, the coumarin core moiety plays a hub role in connecting three domains by forming aromatic contact with Phe422 (Domain 2), hydrogen bond interaction with nitrogen atom from Val147 and Ser148 (Domain 1), and hydrophobic interaction with side chains of Val147 (Domain 1), Gln753 (Domain 4). The hydroxyl benzoate isopentyl group forms hydrophobic contact with several residues, including Pro145, Phe146, Met220, and Asp223, while the novobiose sugar mainly mediates interactions through forming hydrogen bonds with the side chain of Lys151 and Thr175, and week polar interactions with Glu423, Gln749, and Ser750 (Figure 2.a,b).

**Fig. 2.**
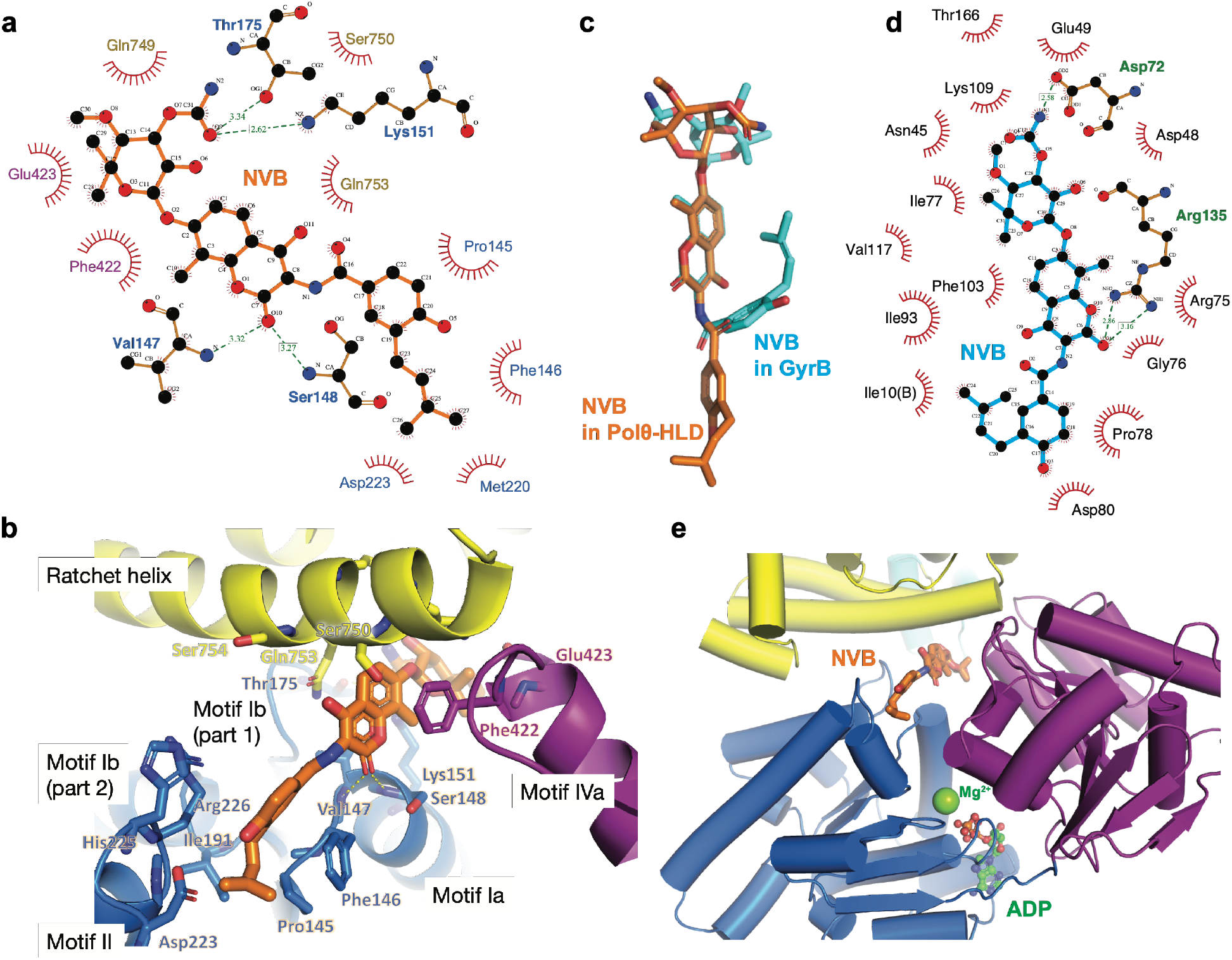
The inhibitor binding site in the Polθ-HLD. (a). The binding site of NVB in the Polθ-HLD analyzed with LigPlot^+^ software, residues from domains 1, 2, and 4 are colored with blue, purple, and yellow, respectively. Hydrogen bonds are highlighted with a dashed line. (b). The NVB binding pocket in the Polθ-HLD. The NVB is shown as brown sticks, and domains 1, 2, and 4 are colored blue, purple, and yellow, respectively. (c) The comparison of NVB in the extended conformation (Polθ-HLD) and bend conformation (GyrB, PDB ID: 1KIJ). (d) The binding site of NVB in the GyrB (PDB ID: 1KIJ). (e) The NVB binding site is away from the classical ATP binding pocket (ADP model from PDB ID: 5a9f).

The three entities of NVB: the hydroxyl benzoate isopentyl group, the coumarin core, and the novobiose sugar exhibits an extended conformation, which is different from canonical bend conformation in the ATP binding site (Figure 2.c). In the bend conformation of NVB revealed by several structural studies^28–31^, the hydroxyl benzoate isopentyl group folds back away from the solvent onto the coumarin ring. Remarkably, besides the extensive hydrophobic interactions, two hydrogen bonds on coumarin core and novobiose sugar are quite similar between NVB and GyrB (Figure 2.d). Figure 2.e shows the distinction between non-canonical NVB binding site and ATP binding pocket (ADP model from PDB ID: 5a9f), located in the interface of domain 1 and domain 2^12^. Due to the slight flexibility of the hydroxyl benzoate isopentyl group and the novobiose moiety in the pocket, they show a worse density in the other protomer compared with the coumarin core. Because of the peculiar physicochemical features, the coumarin moiety represents a privileged scaffold for bioactive compound^37^. The structural information of NVB-Polθ-HLD will provide a structural framework for compound optimization. Therefore, it is possible to develop some coumarin derivatives that exhibit higher affinity and better selectivity targeting the Polθ-HLD by the computational approach of structural-based drug design.

### The inhibitory mechanism by blocking DNA binding

Due to the sequence feature of Polθ-HLD being closely related to HELQ/Hel308-type and RecQ-type helicases, it is not surprising that Polθ-HLD exhibits both DNA unwinding and annealing activities like them^38^. Purified Polθ-HLD can unwind several types of DNA substrates with 3′–5′ polarity, including replication forks^7^, blunt-ended DNA, and DNA with 3′ or 5′ overhangs^39^, but preferentially unwinds DNA with 3’ overhangs. Meanwhile, the Polθ-HLD was also proved to exhibit DNA annealing activity^11^, which it promotes ssDNA annealing in an ATP-independent manner. Polθ-HLD can bind to ssDNA and exhibit bidirectional scanning on the ssDNA^40^. The central tunnel of Polθ-HLD formed by multiple domains represents a putative pathway that ssDNA translocated. As demonstrated in the complex of HEL308 with a partially unwound DNA-duplex substrate, D2 binds the DNA duplex by the extensive positively charged region near the central tunnel and melts the DNA-duplex by a β-hairpin loop. The unwound 3’ tail ssDNA paths through the central tunnel and subsequently interacted with residues from all five domains at different base positions^20^, which provides valuable insights into the ssDNA binding mode and the duplex-unwinding mechanism for this family^20^. Since the first step in a simplified TMEJ model is the recognition and binding of Polθ-HLD to ssDNA tail^6^, an ideal inhibitor can likely block the initial ssDNA recognition step by competing for the protein-DNA interaction.

To gain insight into the inhibitory mechanism by NVB, we superpose the inhibitor-bounded Polθ-HLD structure to the homologous Hel308-DNA complex^20^ (Figure 3.a). Although the overall structures of Polθ-HLD and Hel308 are similar, differential conformational changes happen in different domains. The r.m.s.d values of individually superposing five domains (D1-D5, 1.3, 1.1, 1.1, 3.0, and 0.9, respectively) indicate that the D4 exhibit the most extensive global conformational changes. It’s reasonable when considering the fact that D4 coordinates nucleotide base moiety with D1 and D2 and provides an ideal ratchet for the bidirectional progression of the ssDNA tail, energizing by ATP hydrolysis in an inchworm-like transport mechanism^20^. Meanwhile, the movement of each domain in the DNA-bound modeling state enables a larger space in the central tunnel, which implies conformational changes when ssDNA and DNA duplex binds to the Polθ-HLD. Notably, in the modeling complex, the NVB overlap with 3’ overhang ssDNA (Figure 3.a). In the Polθ-HLD-NVB complex, the NVB vertically wedges between the overhang ssDNA’s third and fourth unpaired bases (Figure 3.b). Amino acids from motif Ia (Val147, Ser148, and Lys151) and motif 1b (part1) (Thr175) in Polθ-HLD interact with the different parts of NVB moiety by hydrogen bond interaction, while the equivalent residues at motif Ia and 1b (part1) in Hel308 directly interact with the DNA backbone of the third and fourth unpaired bases. Therefore, the NVB inhibitor competitively occupied the DNA translocation pathway resulting in the blocking of initial DNA recognition. Considering the activity difference of ssDNA (EC50 about 300nM) and NVB (IC50 about 14uM) to Polθ-HLD, respectively, the NVB at micromolar level affinity is not an ideal candidate inhibitor for clinical treatment. Further structural-based optimization may enable the discovery of low nanomolar potency derivatives with better target selectivity.

**Fig. 3.**
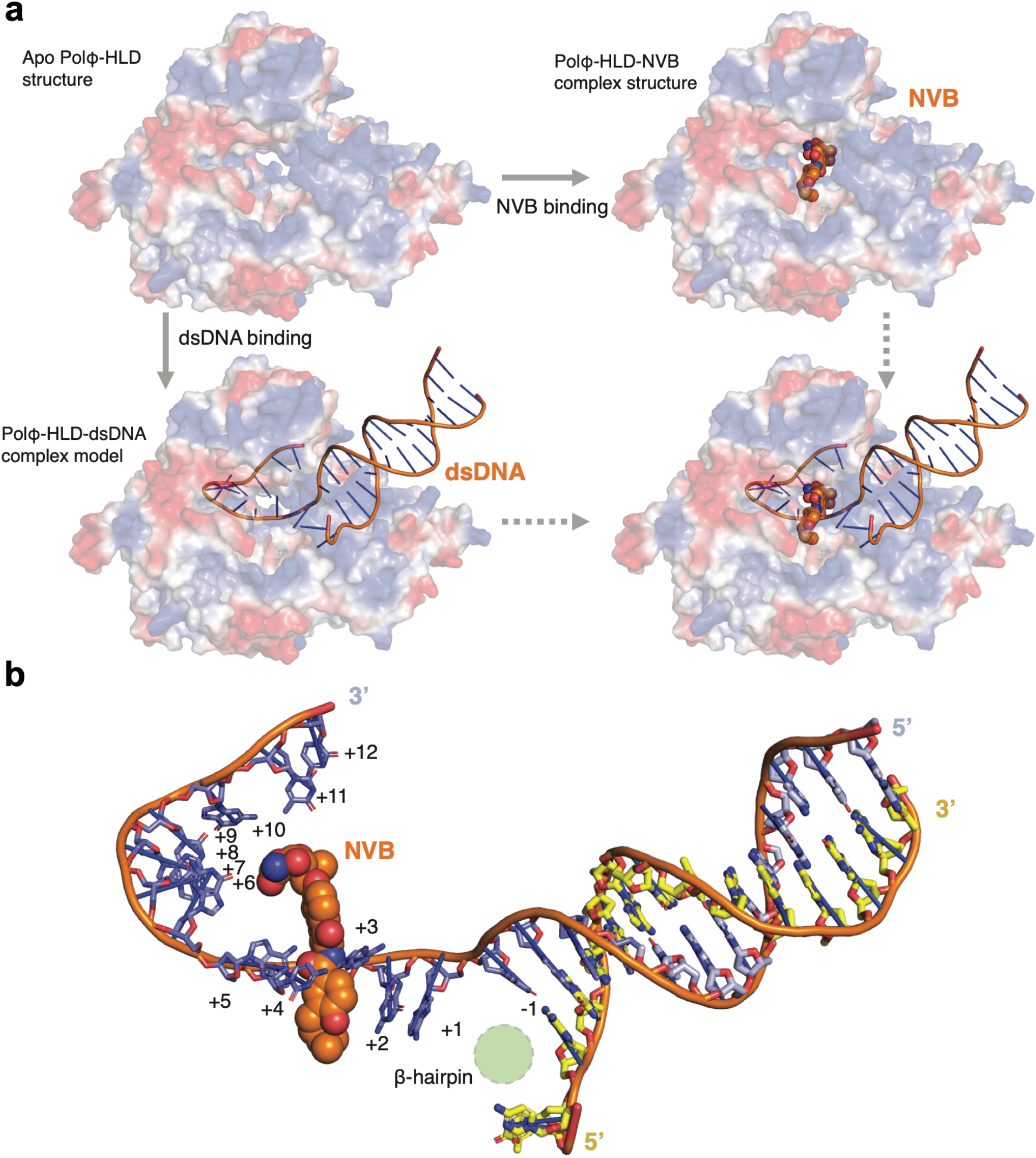
The inhibitory mechanism of NVB. (a). Possible models of NVB and ssDNA binding Polθ-HLD are constructed by homolog modeling with the Hel308-DNA complex (PDB ID: 2P6R). Overlaying the inhibitor binding structure and dsDNA complex model indicates that they directly compete. (b). A detailed structural model of the inhibition mechanism. The NVB binding site is closed to the β-hairpin. The NVB vertically wedges between the third and fourth unpaired bases of the overhang ssDNA.

### The homodimer interfaces

Dimerization plays a role in forming DNA synaptic complexes in the DSB repair processes^14,42–44^, including the NHEJ, HR, and TMEJ process. Dimeric proteins or protein complexes can tether DNA end breaks together and facilitating the formation of intermediate DNA synaptic end complexes. Previous *in vitro* study suggests that although the purified Polθ-POL can perform MMEJ on short ssDNA, full-length POLQ is essential for MMEJ on long ssDNA with a dimeric model^14^, in which the Polθ-HLD can not only bridge two DNA end breaks together but also suppress the intrastrand pairing activity of long ssDNA.

Consistent with the previous model, the apo dimeric Polθ-HLD shares a similar dimer interface formed by chain A and chain C in the tetramer Polθ-HLD (PDB: 5A9J), which is mainly contributed by interchain hydrogen bonds and hydrophobic contact from D4. Interestingly, we find that the dimeric interface of apo Polθ-HLD in the Cryo-EM structure is slightly less extensive than in the crystal structure (580 Å^2^ vs. 850 Å^2^ interface area, calculated by the PDBePISA server^45^), which reflects the structural flexibility of dimer complex in solution. In contrast, probably due to the stabilization effect of inhibitors on inter-domain dynamics, the NVB-occupied structure exhibits a larger extent of dimer interface than the apo structure (857 Å^2^). Meanwhile, we also observed some extent of conformational change in D5, resulting in forming an interprotomer hydrogen bond between Arg791 and Asn773. Likewise, the conformational change of D5 was also proposed to modulate the DNA binding or other partner interaction with an autoinhibitory mechanism^25,39^. Because the dimerization of Polθ-HLD will bring the 3’ ssDNA overhangs in close proximity, it was suggested that it could directly catalyze the annealing reaction of complementary DNA^11,19^ or facilitate the DNA annealing activity of Polθ-POL by bidirectional movement as a dimer on the long ssDNA^14^. Although the inhibitor binding site we characterized here is far from the dimerization site, small molecules or antibodies targeting the dimeric interface may exhibit an inhibitory effect on Polθ functionality.

## Discussion

Mammalian cells have evolved multiple pathways to repair DNA double-strand breaks (DSBs) and ensure genome stability. The specific mechanism of how the TMEJ involving has received increasing attention in recent years. Because Polθ is an error-prone polymerase, it repairs the DNA ends by annealing micro-homologous sequences resulting in deletions and insertions in the break sites. As a result, this type of repair can promote genetic instability in the early stage of tumorigenesis to promote cancer progression. Therefore, the Polθ gene is upregulated in numerous cancers, and its overexpression is associated with poor prognosis. Developing Polθ inhibitors represents a promising clinical treatment strategy^46–48^.

Here, we report the inhibitor-occupied structure of Polθ-HLD and characterize a promising allosteric inhibitor binding site distinct from the canonical ATP binding site. Notably, the inhibitor binds to a putative ssDNA translocation pathway, which blocks ssDNA binding. NVB is an antibiotic^49^ that has been withdrawn from the market due to some unfavorable safety profiles and was recently proven to effectively mediate HR-deficient tumor cell death by targeting Polθ-HLD^18^. Strikingly, NVB was historically reported to have an inhibitory effect on DNA repair^50,51^, and it was actually the subject of oncology trials in phase 1^52,53^ and phase 2 studies^54^, combined with high-dose chemotherapy. Moreover, NVB was also used to reverse breast cancer resistance protein-mediated drug resistance^55^. The low potency of NVB to Polθ-HLD will restrict the development of clinical cancer treatment. Although the NVB is well-known for its coumarin scaffold moiety, which usually competitively binds to the ATP binding pocket, the Polθ-HLD-NVB complex structure indicates that NVB allosterically binds to Polθ. Because the coumarin moiety pivots the domain-domain interactions and blocks ssDNA interaction, other NVB analogs that keep the coumarin moiety may exhibit a similar inhibitory effect. Actually, a series of novobiocin analogs have been developed to investigate the antiproliferative activity against several cancer cell lines but were supposed to be selective to target HSP90^56,57^. Still, those compounds’ inhibitory efficacy and selectivity on Polθ-HLD haven’t been reported. Collectively, a structural-based hit optimization strategy will accelerate the discovery of analog compounds with higher affinity, better efficacy, and selectivity for clinical treatment.

In summary, our structure provides a structural insight into how NVB inhibits Polθ-HLD activity. Directly competing ssDNA binding sites represent a straightforward strategy to develop Polθ-HLD inhibitors. Since both Polθ-HLD^18^ and Polθ-POL^16^ inhibitors demonstrate selective toxicity in BRCA1/2-deficient cancer cells, patients harboring cancer-specific alterations in specific genes might benefit from clinical Polθ inhibitor treatment. Despite the revolutionary efficacy of PARP inhibitors for the treatment of BRCA-mutated tumors, the acquisition of PARP inhibitor resistance has been observed in many patients. Therefore, PARP inhibition combined with Polθ inhibition can be a promising synthetic lethal therapeutic strategy to improve cancer treatment.

## Methods

### Expression and purification of the human Polθ-HLD

A truncated version of the human Polθ-HLD (residues 67-940) was cloned into the pFastBac-1 vector with an amino-terminal 8×His tag and a carboxy-terminal Twin-Strep-tag. The protein was expressed in the Sf9 insect cell (Expression system) in the ESF921 medium using the Bac-to-Bac baculovirus system. The purification of the Polθ-HLD protein is mainly referring to the methods described previously with slight modification^19^. Briefly, the Polθ-HLD protein was expressed for 48 hours after infection with recombinant baculovirus, and cells were collected and lysed using a buffer containing 25 mM Tris (pH 8.5), 500 mM NaCl, 0.5 mM Tris (2-carboxyethyl) phosphene (TCEP), 0.0025 mg/mL Leupeptin. Ni-NTA affinity purification was used as the initial purification step and followed by Strep affinity chromatography for further purification. The eluted proteins were concentrated, aliquoted, and flash-frozen for storage at -80°C.

### ATPase activity assay

The ATPase activity assay followed a previously established protocol using the ADP-Glo kinase assay (Promega)^18^. The 30mer ssDNA (5’-CCAGTGAATTGTTGCTCGGTACCTGCTAAC-3’) used in the assay was synthesized from Sangon Biotech. The reaction was done in 20ul condition in 384 optiplate (PerkinElmer), and the reaction buffer contains 40mM Tris-HCl pH 7.5, 20mM MgCl2, 0.01% Triton X-100, 0.01% BSA, and 1mM DTT.

The final concentrations were 30 nM of purified Polθ-HLD protein, 20 μM of ATP, and 600 nM of 30mer ssDNA, with DMSO or NVB. The order for adding the components were: 2.5 μl of 3x NVB or DMSO, 2.5 μl of 3x POLθ (incubate at room temp for 15 min), and finally, 2.5 μl of 3x mixture containing ATP and ssDNA, and all components were prepared in 1x reaction buffer. DMSO wells represented 0% inhibition, while no-enzyme wells represented 100% inhibition. Plates were covered with an aluminum seal and incubated at room temperature. 2 hours later, 7.5 μl of ADP-Glo reagent (Promega kit) was added to each reaction well and incubated at room temp for 40 min. Next, 15 μl of Kinase detection reagent (Promega kit) was added to the wells, and plates were incubated for 1 hour. Finally, ATP hydrolysis was quantified by luminescence measured on the Ensight multimode plate reader (PerkinElmer), and data was analyzed with Prism 8 software (GraphPad Prism).

### Cryo-EM sample preparation and data collection

#### Cryo-EM Sample Preparation

For the preparation of the inhibitors complex, the purified Polθ-HLD was mixed with a 50-fold molar excess of inhibitor NVB, and incubated on ice for 30 min before applying for grid preparation.

The purified apo Polθ-HLD at a protein concentration of 1mg/mL was applied for grid preparation on glow-discharged holey gold grids (Quantifoil Au R1.2/1.3, 400mesh) and GraFuture™ GO grid (Quantifoil Au R1.2/1.3, 400mesh). An aliquot of 4 μL protein sample of Polθ-HLD-NVB complex at a protein concentration of 0.92 mg/mL was loaded onto a glow-discharged 400 mesh grid (Quantifoil Au R1.2/1.3), blotted with filter paper for 3.0 s and 3 blot force, and then plunge-frozen in liquid ethane using a Thermo Fisher Vitrobot Mark IV.

#### Data Collection

Cryo-EM micrographs were collected on a 300kV Thermo Fisher Titan Krios G3i electron microscope equipped with a K3 direct detection camera and a BioContinuum energy filter (GIF: a slit width of 20eV). The micrographs were collected at a calibrated magnification of x105,000, yielding a pixel size of 0.3345 Å at a super-resolution mode. In total, 1,824 micrographs were collected at an accumulated electron dose of 50e^-^Å^-2^ s^-1^ on each micrograph that was fractionated into a stack of 32 frames with a defocus range of -1.0 μm to −2.0 μm.

### Cryo-EM data processing, model building, and refinement

#### Image processing

Beam-induced motion correction was performed on the stack of frames using MotionCorr2 ^53^. The contrast transfer function (CTF) parameters were determined by CTFFIND4 ^54^. A total 1,824 good micrographs were selected for further data processing using cryoSPARC^55^. These micrographs were then curated to remove suboptimal data, leaving 1,599 micrographs. 1,311,914 Particles were auto-picked by the blob picker and template picker program in cryoSPARC. After 2 rounds of 2D classification, 655,791 particles were selected from good 2D classes and were subjected to *ab-initio* reconstruction, followed by heterogeneous refinement. Further homogeneous refinement and non-uniform refinement were conducted for 311,924 particles from the best 3D classes without applying symmetry, which resulted in a 3.14 Å map for the Polθ -HLD-NVB complex protein based on the gold-standard Fourier shell correlation criterion at FSC=0.143. The local resolution was then calculated on the final density map.

#### Model building and refinement

The model of Polθ-HLD-NVB complex was built by fitting a structure of the complex (predicted by AlphaFold2) into the density map using UCSF Chimera^56-57^, followed by a manual model building of the complex molecules in COOT^58^ and a real space refinement in PHENIX^59^. The model statistics were listed in Supplementary Table 1.

## Contributions

H.B.G and Y.T.H performed protein expression and purification experiments, Y.X.W and H.M.Z. prepared grid, collected data, and processed data, J.M. performed functional assay, Y.X.W,H.M.Z, Y.D.H, and J.L. built and refine the model, and assisted structure analysis, Y.J.L, G.Z., X.D.N, F.Y.Z., J.H, and A.G supervised the project experiments and analyzed the data, X.D.N, F.Y.Z., J.H. wrote the manuscript.

## Supplementary Figures

**Fig S1.**
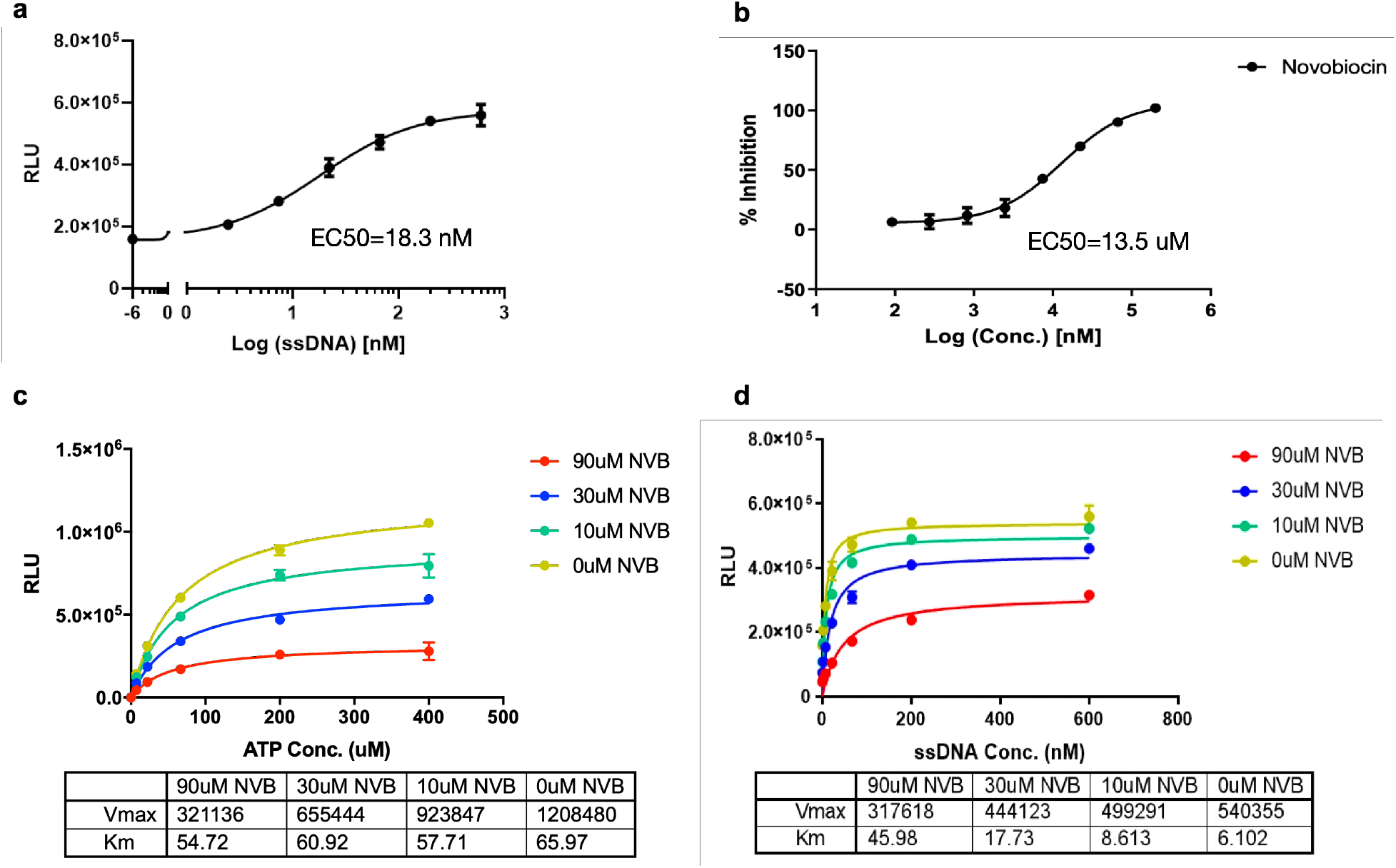
The functional validation of NVB inhibitory effect on ATPase activity. (a). The ssDNA stimulates the ATPase activity of Polθ-HLD with an EC50 of about 18.3 nM. (b). The dose-response curve of NVB inhibits ATPase activity with EC50 of about 13 uM. The maximized inhibition at 200uM NVB was normalized to 100% inhibition. (c). The ATP competition assay. The NVB exhibits non-competitive inhibition with respect to ATP. (d) The ssDNA competition assay. The NVB exhibits competitive inhibition with respect to ssDNA. Each data point represents the mean ± SD from two technical replicates.

**Fig S2.**
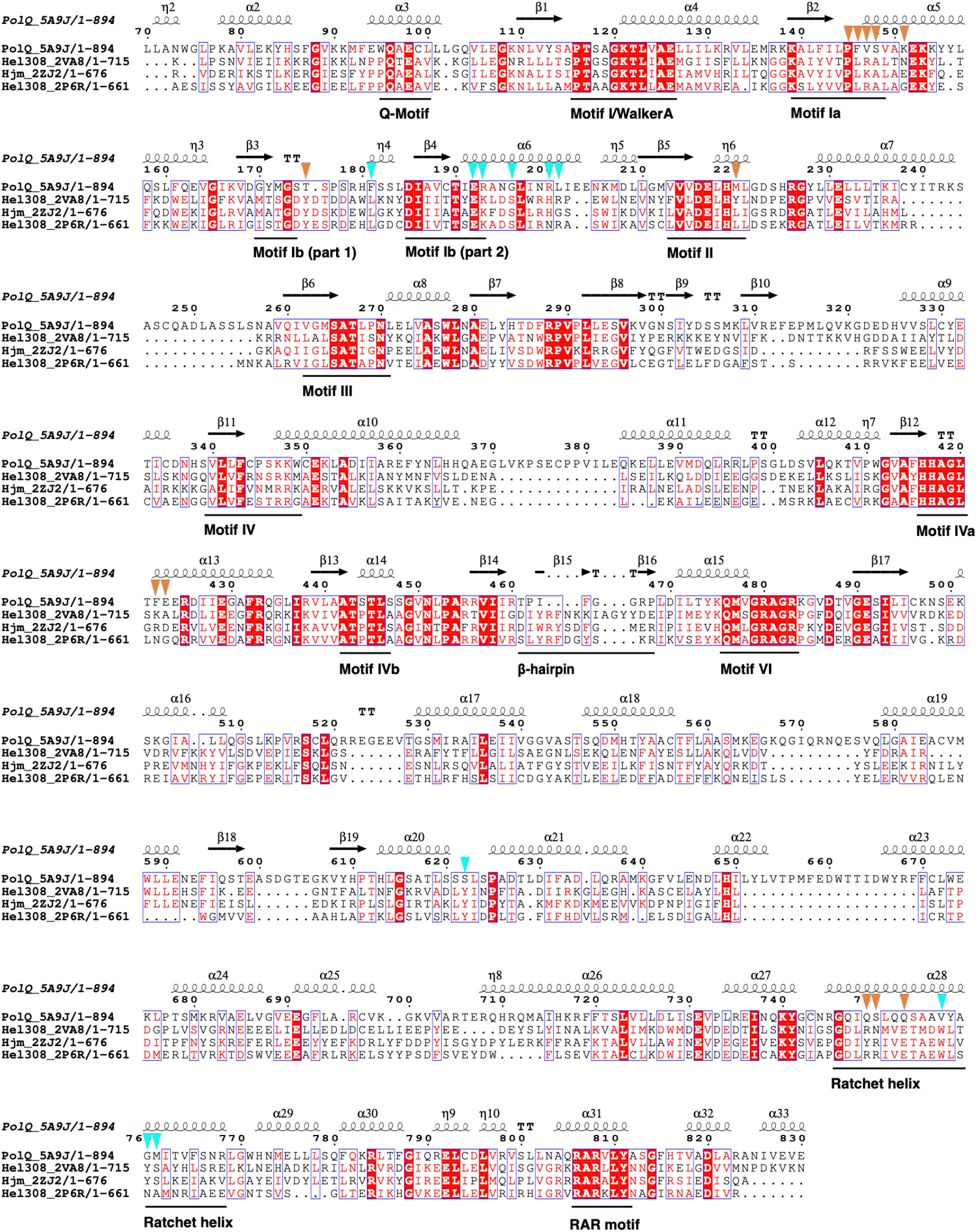
The sequence alignment and conserved motifs of four superfamily 2 helicases with crystal structures. The selected regions of Polθ-HLD (residues 70-830) were annotated, and the medium-conserved residues of those four proteins are shown in red text, and the high-conserved residues are shown in white text with red background. The secondary structure of Polθ-HLD on top of the alignment was automatically generated with ESPript 3.0 server. The residues that interact with NVB and 236 were noted as brown and cyan triangles, respectively.

**Fig S3.**
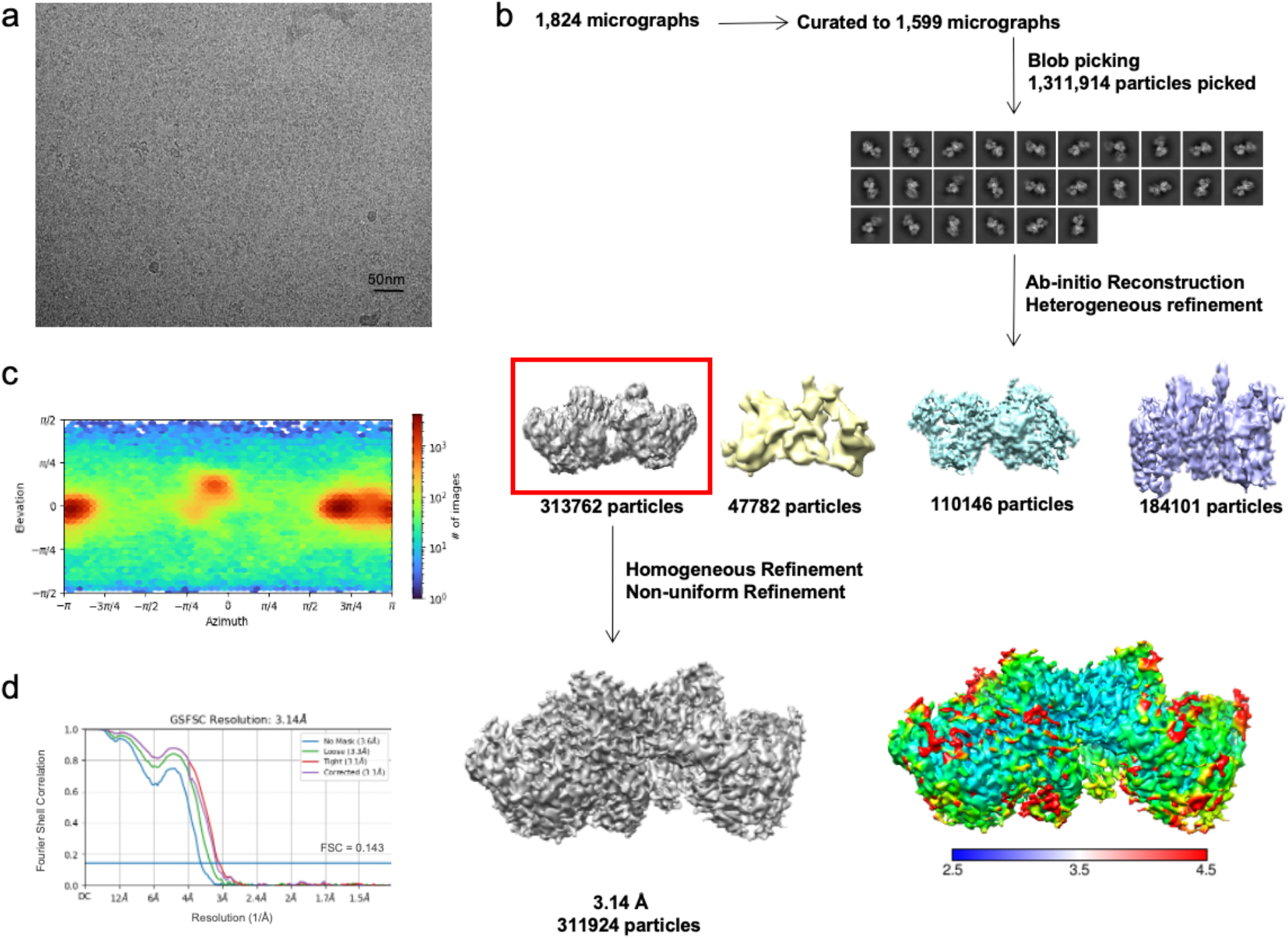
Cryo-EM analysis of the Polθ-HLD. (a). The micrograph of Polθ-HLD. (b). Flow chart for cryo-EM data processing of the Polθ-HLD. (c). Direction distribution of particles in the final Polθ-HLD refinement. (d). FSC curves of the final Polθ-HLD map and model validation.

